# Relating evolutionary selection and mutant clonal dynamics in normal epithelia

**DOI:** 10.1101/480756

**Authors:** Michael W J Hall, Philip H Jones, Benjamin A Hall

## Abstract

Cancer develops from mutated cells in normal tissues. Whether somatic mutations alter normal cell dynamics is key to understanding cancer risk and guiding interventions to reduce it. An analysis of the first incomplete moment of size distributions of clones carrying cancer-associated mutations in normal human eyelid skin gives a good fit with neutral drift, arguing mutations do not affect cell fate. However, this suggestion conflicts with genetic evidence in the same dataset that argues for strong positive selection of a subset of mutations. This implies cells carrying these mutations have a competitive advantage over normal cells, leading to large clonal expansions within the tissue. In normal epithelium, clone growth is constrained by the limited size of the proliferating compartment and competition with surrounding cells. We show that if these factors are taken into account, the first incomplete moment of the clone size distribution is unable to exclude non-neutral behavior. Furthermore, experimental factors can make a non-neutral clone size distribution appear neutral. We validate these principles with a new experimental data set showing that when experiments are appropriately designed, the first incomplete moment can be a useful indicator of non-neutral competition. Finally, we discuss the complex relationship between mutant clone sizes and genetic selection.

**Significance Statement:** Aging normal epithelial tissues are extensively colonized by clones carrying cancer associated mutations. Insight into the emergence of mutant clones is key to guide cancer prevention. However, the statistical evidence as to whether mutant clones emerge by neutral competition or due to a competitive advantage conferred by mutation is conflicted. We reconcile this apparent contradiction by demonstrating that the previously presented metrics for measuring neutrality from clone sizes are dependent on the spatial constraints imposed by the tissue structure and experimental design. Furthermore, we show that clonal competition within a recently reported dataset of healthy human esophageal tissue is non-neutral. Finally, we discuss how discrepancies between measures of clone size and genetic selection can provide insight into early carcinogenesis.

## Introduction

Large scale sequencing of cancer genomes has led to the discovery of many recurrently occurring genetic mutations that are potential “drivers” of the disease (1–3). Recently, however, a number of studies investigating normal tissues have found that many of these mutations are also present in apparently healthy tissue (4–9). To understand tumorigenesis it is therefore important to study the acquisition and spread of mutations in normal tissue.

The commonest human cancers are derived from squamous epithelia, which consist of layers of keratinocytes (10). Cells are continually shed from the tissue surface and replaced by proliferation (**Fig. 1a**). The proliferating cells accumulate mutations over time (11). If such clones persist within the otherwise normal tissue, they may acquire the additional genomic alterations that lead to cancer (11). A key question is whether these large, persistent clones arise by neutral competition or are a consequence of cancer-associated mutations increasing the competitive fitness of mutant cells above that of wild type cells. If the former, little can be done to alter the risk of cancers emerging from mutant clones in normal tissue. However, if the founding clones of cancers emerge by competitive selection, it is possible that interventions that alter the fitness of mutant cells may decrease cancer risk. A dataset of mutant clones detected in normal human eyelid skin appears to contain conflicting evidence supporting both neutral and non-neutral mutant cell dynamics (4, 12). This “paradox” has yet to be resolved (13, 14), leading to uncertainty over the somatic mutant cell dynamics in normal epithelial tissues (15). This paper aims to unpick this apparent inconsistency.

**Figure 1.**
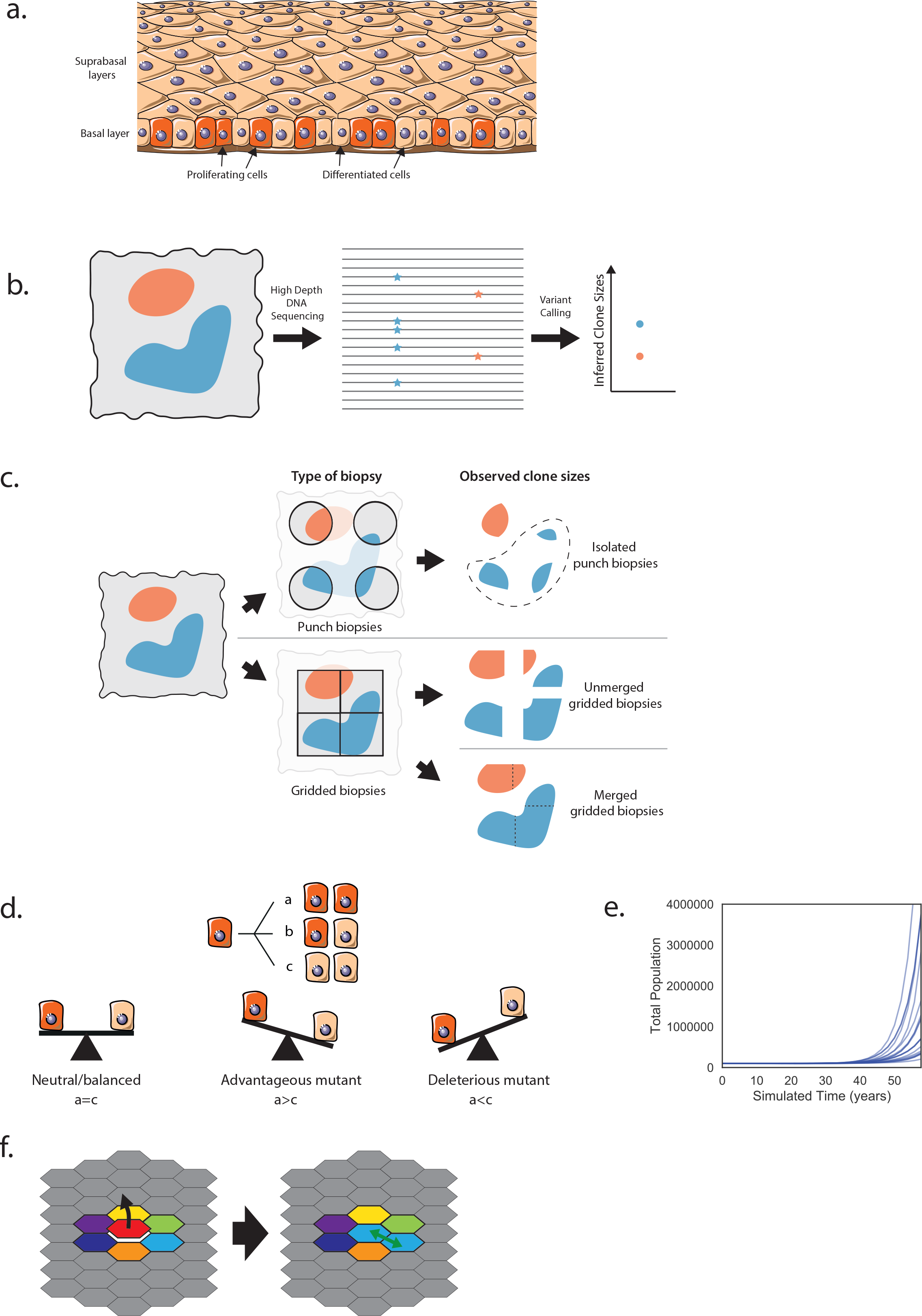
Data collection and cell dynamics. **a)** Proliferation occurs in the basal layer of the epithelium. After differentiation, cells migrate through the suprabasal layers before being shed. Image from smart.servier.com, licensed under CC BY 3.0, edited from original. **b)** DNA from a biopsy (left) containing mutant clones (red and blue) is sequenced and the variant allele fraction (proportion of reads containing the mutation, middle) used to infer clone sizes (right). **c)** The method of taking biopsies can affect the observed mutant clone size distribution. Isolated punch biopsies (top) may not capture the entirety of a mutant clone; in the analysis in (12), clones which spanned multiple biopsies (shown in dashed area) were excluded. Ungapped gridded biopsies (bottom) enable the reconstruction of larger clone sizes. **d)** The stochastic single progenitor model of cell dynamics. Each dividing cell (red) can produce two dividing cells (a), two non-dividing differentiated cells (brown) (c), or one of each type (b). In a homeostatic tissue or neutral clone, the probabilities of each symmetric division option are balanced (left). An advantageous mutation would increase the proportion of dividing cells produced (middle), and a deleterious mutation would increase the proportion of differentiated cells (right). Note that in the non-neutral case, the probabilities of each division type do not have to be fixed over time, but can depend on the cell context. **e)** Simulation of the model shown in **d)**. If mutations introduce perpetual positive fate imbalances then the population will eventually explode. Total population of 20 simulations with mutations introducing only small fate imbalances drawn from N~N(mean = 0.25%, std = 1.25%). **f)** In the spatial Moran process, a differentiating cell (red) is replaced by the division of a neighboring cell (light blue).

The dataset in question is from a study of mutations in normal human eyelid skin epidermis (4). DNA was extracted from small samples of epidermis. About 500 DNA molecules of each targeted gene were sequenced and compared to the genome of the same tissue donor (**Fig. 1b**). Somatic mutations were detected as altered sequences present in one or more samples (4). The proportion of altered DNA reads containing a mutation, the variant allele fraction (VAF), was assumed to be proportional to the size of the mutant clone (**Fig. 1b**).

An analysis of inferred sizes of the mutant clones in the human eyelid data argues for neutral dynamics (12). Observed clone sizes can be compared to the predictions of candidate mathematical models of cell dynamics to determine the best-fitting model (16). Lineage tracing experiments in homeostatic, unmutated mouse epidermis and esophagus suggest that these tissues are maintained by a single population of equipotent progenitor cells (**Fig. 1d**) (17–19). The outcome of individual cell divisions is unpredictable but on average 50% of the progeny of dividing cells differentiate, exit the proliferative compartment and are eventually shed from the tissue while 50% remain to divide again. Such balanced stochastic cell fate leads to wide variation in clone sizes while the total cell population remains constant. Mutant clone sizes observed in the human eyelid skin have been compared to predictions from this neutral stochastic model (12). The comparison is made using the first incomplete moment of the clone size distribution (**methods**), which has been used in several studies to shed light on mutant clone growth dynamics (12, 20–23). In economics the first incomplete moment is used to study inequality - it shows how much of the wealth is held those with an income higher than *x* (24). It has a similar role for the clone size analysis - it shows the proportion of the mutated cells that are in clones of size *x* or larger. Using the first incomplete moment has two advantages over using the clone size distribution directly. Firstly, the first incomplete moment reduces the fluctuations caused by low sample size (20) and secondly it simplifies the comparison of data to the neutral theory. The neutral model predicts that the first incomplete moment of mutant clone sizes will have a negative exponential form (12) - where there are many small clones and few large clones. A deviation from the exponential shape could indicate non-neutral competition - some clones have expanded to take over more than their expected share of the tissue. To make it easier to see this deviation, we use the logarithm of the first incomplete moment (LFIM), which turns the exponential curve into a straight line. Non-neutral competition would then be indicated by a deviation from the straight-line (12). Using this criterion, the inferred mutant clone sizes from the human eyelid appear largely consistent with the neutral model (12).

However, the theory of neutral competition of cancer-associated mutations is incompatible with results from several mouse and human studies that observed non-neutral mutant expansions in normal epithelial tissues (18, 20, 25). Furthermore, signs of non-neutral clonal competition in the eyelid mutational data can be detected using dN/dS analysis, a method from population genetics. This examines the ratio of protein-altering mutations (dN) to silent mutations (dS) for each gene (26). Once relevant corrections have been applied, a dN/dS ratio of 1 is indicative of neutral behavior. A dN/dS value of less than 1 indicates the mutated gene has a negative effect on the competitive fitness of mutant cells compared with normal cells. However, if a disruption of the protein provides a growth advantage to the cell, then the number of protein-altering mutations that expand to a clone of detectable size will be increased, leading to a dN/dS ratio greater than 1. Analysis of the human eyelid mutations found 6 of the 74 sequenced genes had significantly raised dN/dS ratios ranging from 3 to over 30, consistent with mutations in those genes driving clonal expansion (4). Additionally, protein-altering missense mutations in some driver genes, e.g. *NOTCH1*, *NOTCH2* and *TP53*, were not randomly distributed but concentrated in functional domains. This suggests positive selection of function-altering mutations, which is also incompatible with neutrality (4).

Here we show that in some conditions *non-neutral* competition can produce a straight-line LFIM, and therefore a straight LFIM alone is not a clear indicator of *neutral* dynamics. We can thus reconcile clone size distributions and positive genetic selection.

## Results

We began by noting several important constraints that apply to the mutant clones in normal epithelia. Firstly, the cellular structure and composition of the tissue remains at least approximately constant. Secondly, the proliferative compartments of the epidermis and esophageal epithelium contain few barriers to mutant clone expansion. In a tissue with a high burden of mutations like the human eyelid, this means that expanding clones will soon collide and compete with each other as well as with un-mutated cells. These two constraints were not included in the mathematical model used in the first incomplete moment analysis of the human eyelid data (12), meaning mutant clones in this model with a growth advantage (**Fig. 1d**, middle) could expand without limit (**Fig. 1e**).

### Spatial constraints alter clone size distributions of non-neutral mutations

To address the effect of clonal competition we used a mathematical model drawn from the study of population genetics. We ran Moran-type (27) simulations of cell competition on a 2D grid to represent the epidermis (**Fig. 1f**). Cells lost through differentiation are replaced by the division of a neighboring cell (**methods**), similar to behavior observed in mouse epidermis (28). During each division, there is a small chance that one of the daughter cells will acquire a mutation (**methods**).

Simulations of a 2D neutral model produced an approximately straight LFIM (**Fig. 2a**). A non-neutral spatial model in which a small proportion of mutations change the fitness of a cell (**methods**) may deviate from a neutral appearance by curving away from the straight line (**Fig. 2b**). This was due to the contrast between the relatively large non-neutral clones and the smaller clones growing neutrally. Surprisingly, however, simulations with a higher proportion of non-neutral mutations may generate a straight line (**Fig. 2c**). This is because almost all of the simulated tissue is taken over by non-neutral clones (**Fig. 2d**). The only neutral mutations that persist are those that occur in clones with non-neutral mutations, carried as ‘passengers’. As all remaining clones exhibit similar behavior, the LFIM is straightened. This shows that a straight-line LFIM does not necessarily imply neutral competition and is consistent with positive dN/dS ratios (**Fig. 2e**). We concluded that since there is a high burden of mutant clones in the eyelid, the tissue is likely to be extensively colonized by non-neutral mutant cells, contributing to the apparently neutral appearance of the clone size distribution.

**Figure 2.**
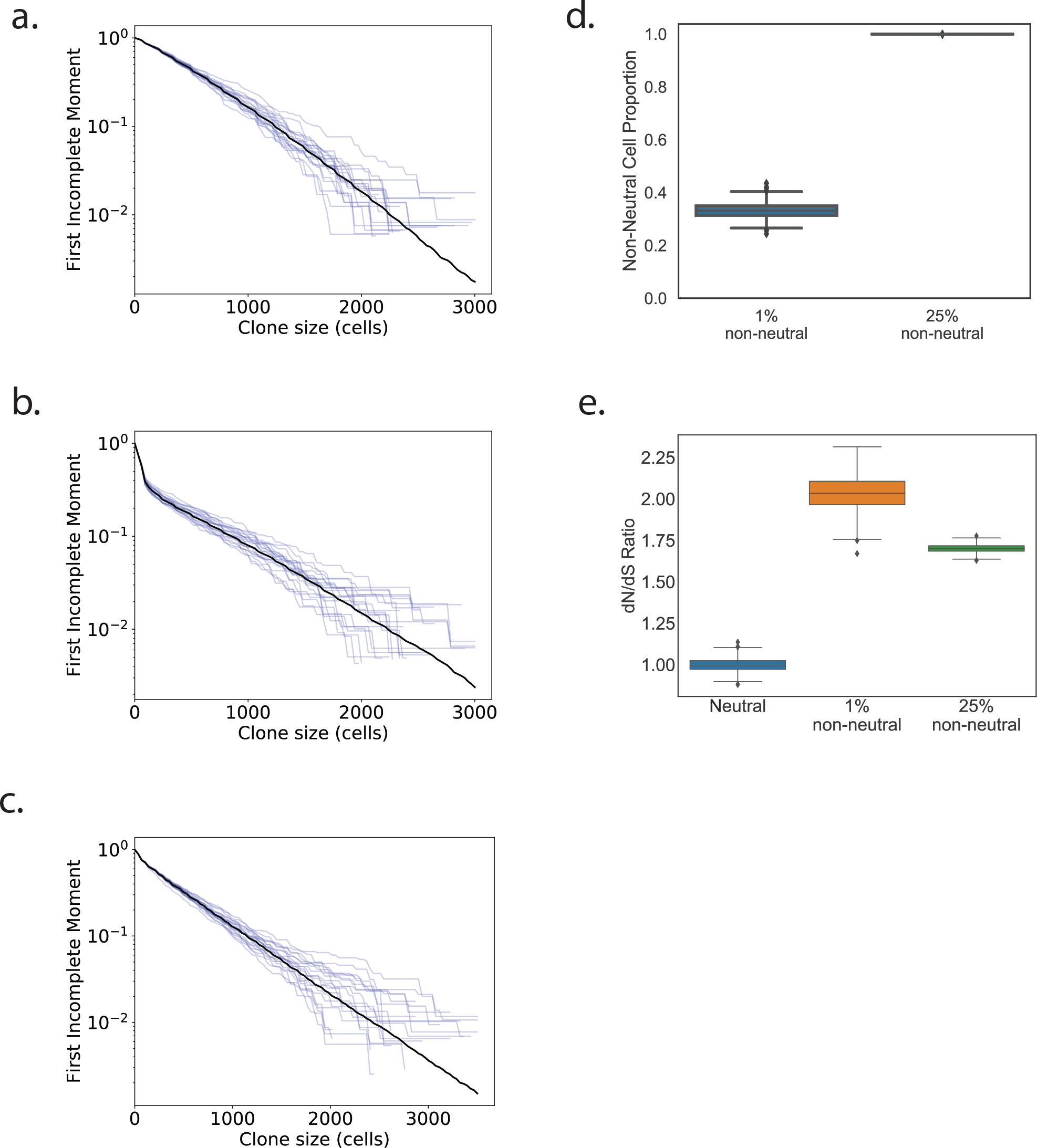
First incomplete moments of 2D simulations. **a-c)** First incomplete moments of 2D simulations (**methods**). The average of 1000 simulations is shown in black, a selection of 20 individual simulations is shown in blue. **a)** Neutral simulations. **b)** Simulations where 1% of mutations are non-neutral. A deviation from the straight line is seen at clone sizes of approximately 100 cells. **c)** Simulations where 25% of mutations are non-neutral. **d)** Proportion of cells at the end of the simulations with a fitness altered by non-neutral mutations. In the 25% non-neutral simulations, by the end of the simulation almost the entirety of the tissue has been colonized by non-neutral mutant clones. **e)** dN/dS values from the simulations shown in **a)-c)**. To enable this calculation for the neutral simulations, a proportion of neutral mutations were labelled as non-neutral but did not affect cell fitness.

### Impact of sampling methods on measurement of clone size distributions

Another consideration that may impact the measurement of clonal size distributions and hence inference of mutant clone dynamics is experimental design. In the human eyelid study, spatially separated tissue samples were collected (**Figure 1c**) (4). The area of the sample defines an upper limit on the reliable estimation of clone size. In the eyelid experiment, the area of each sample was less than that of the largest clones. The lower limit of clone size detection is also related to the sample area, since mutations present in only a small fraction of the cells in the sample may not be detected due to the technical noise in DNA sequencing (29). We simulated the combined effects of spaced samples in which only clones occupying 1% or more of the area of the sample can be detected (**methods**). **Figures 3a,b,c** show these effects on the first incomplete moments of the simulations from **Figures 2a,b,c** respectively. The results lead us to conclude that these experimental factors may artefactually reduce a deviation of LFIM from a straight line caused by non-neutral competition (**Figures 2b**, **3b**).

**Figure 3.**
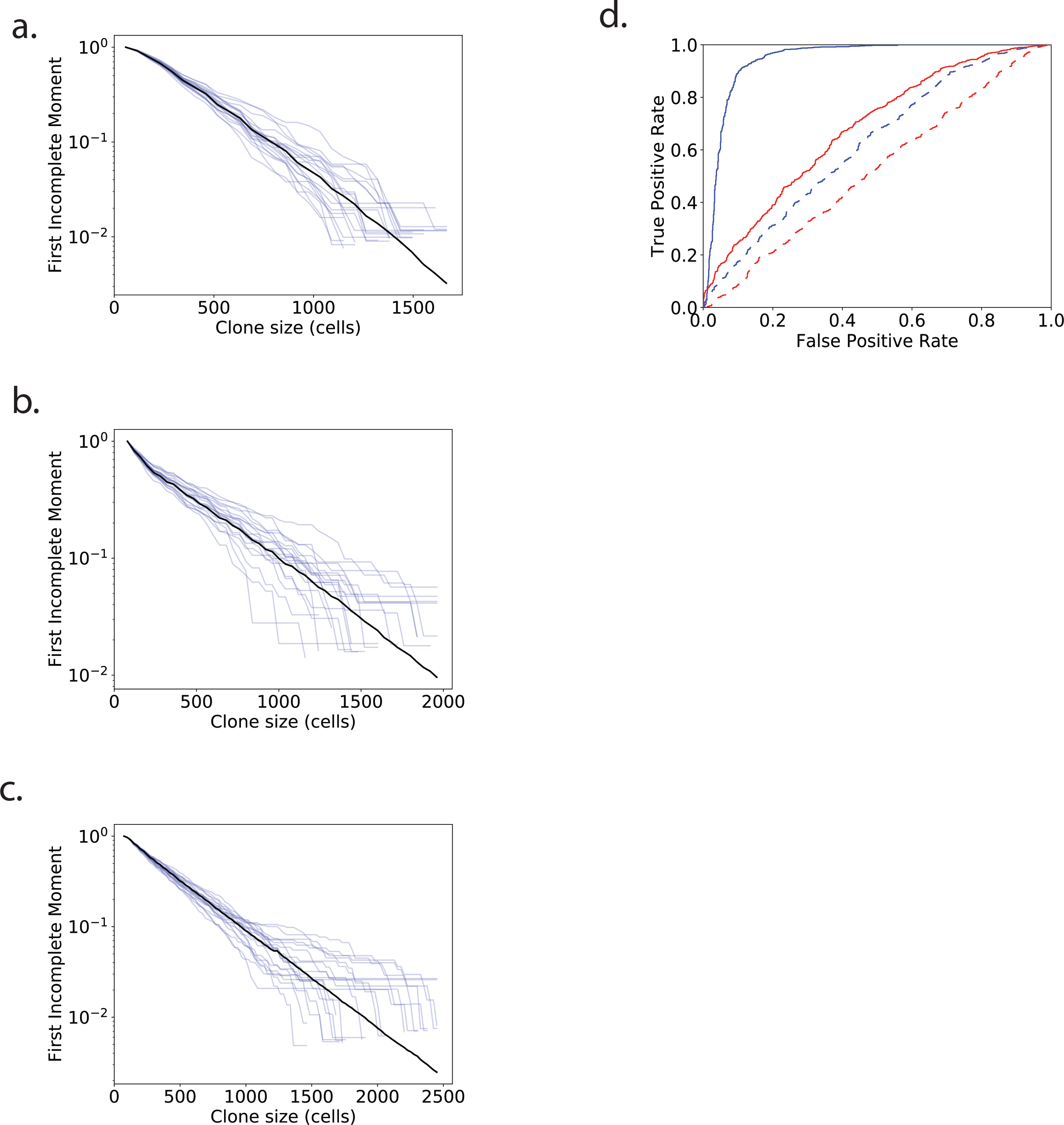
First incomplete moments of 2D simulations with biopsy sequencing. **a-c)** The simulations from **Figure 2a,b,c** respectively, with the effects of biopsy and sequencing. **d)** ROC curves using R^2^ of the log first incomplete moment of the clone size distribution as the classifying statistic. Red, simulated biopsy and sequencing; blue, full data randomly subsampled to match biopsy plus sequencing sample sizes. Solid, 1% non-neutral; dash, 25% non-neutral. Area Under the Curve (AUC) is a measure of how successful the classifier is at distinguishing the two groups. A perfect classifier will have an AUC of 1. A random guess will have an AUC of 0.5. AUCs: Full data, 1% non-neutral, 0.94; biopsy plus sequencing, 1% non-neutral, 0.68; full data, 25% non-neutral, 0.62; biopsy plus sequencing, 25% non-neutral, 0.52.

### Ability of LFIM to resolve neutral competition versus selection

We next tested how well the LFIM could discriminate between the neutral and non-neutral simulations using the coefficient of determination, R^2^, to measure the straightness of a line, as in previous studies (12, 23) (**methods**). For the LFIM to be a successful indicator of neutrality, the neutral simulations need to have a higher R^2^ than the non-neutral simulations. Receiver Operating Characteristic (ROC) curves show the accuracy of the LFIM as a test of neutrality depended on both the underlying shape of a non-neutral clone size distribution and on the experimental sampling method (**Fig. 3d**). For example, using the LFIM was little better than a random guess (Area under the curve, AUC = 0.52) when attempting to distinguish the neutral simulations from **Fig. 3a** from the non-neutral simulations in **Fig. 3c**, where the underlying shape of the non-neutral LFIM was largely straight (**Fig. 2c**) and the clones were measured using simulated DNA sequencing of spatially separated biopsies.

### Human esophageal mutant clone sizes demonstrate non-neutral growth

To validate this analysis we drew on a second experiment that measured mutant clones in normal human esophageal epithelium (5). dN/dS analysis revealed mutations in 14 of 74 genes sequenced were under significant positive selection (5). This study used an ungapped sampling strategy in which the epithelium was cut into gridded arrays of samples which were then deep DNA sequenced, allowing the areas of clones that extend over multiple samples to be determined (5) (**Figure 1c**). This key difference in design from the eyelid experiment allowed us to investigate the effect of sampling on the incomplete moment analysis of the eyelid data by comparing the gridded data with what would have happened if the esophagus was sampled in the same manner as the eyelid skin (**Fig. 1c**). The LFIMs for clone sizes estimated from both sampling approaches are shown in **Figure 4**. Taking **Figure 4d** as an example, the gapped sampling approach results in an LFIM that fits well with a straight line (R^2^ = 0.96) and therefore appears consistent with neutral competition. However, if using the clone sizes based on a gridded approach, the LFIM deviates from the straight-line (R^2^ = 0.78), suggesting non-neutral competition may have occurred in the tissue. For each of the 9 individuals in the study, the LFIM exhibits a greater deviation from the straight line when using gridded samples than when using spaced samples.

**Figure 4.**
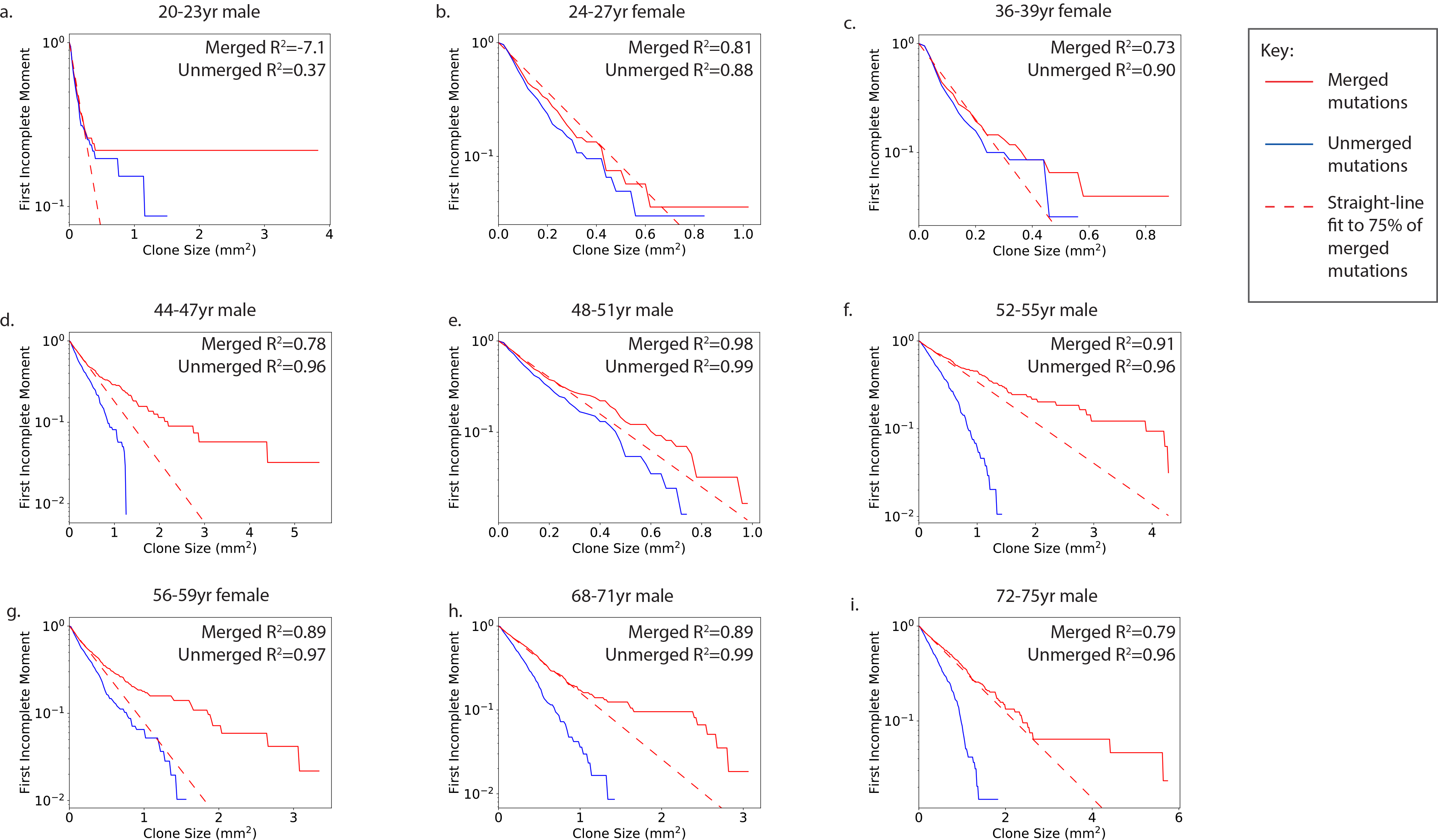
Normal human esophagus. First incomplete moments of the human esophagus mutation data for the nine individuals in the study (5). The clone sizes are either inferred from each 2mm^2^ sample without merging (blue) or by using the gridded system to infer the size of mutations which span multiple samples using the methods of the original study (5) (red, solid). The extent of deviation from the straight line can be seen by comparing the data (solid) to the dashed red line, which shows a straight-line fit to the smallest 75% of clones in the merged case. LOH copy number changes were frequently found to co-occur with protein-altering *NOTCH1* mutations (5) and to obtain conservative estimates of clone sizes we assume this is the case for all protein-altering *NOTCH1* mutations. All other mutations on chromosomes 1-22 were assumed to be heterozygous. R^2^ values can be negative because the line fitting is constrained to pass through the point (*m*, 1), where *m* was the smallest observed clone size. Ages given as a range for anonymization purposes.

An intriguing pattern can be seen within the esophageal data. With the exception of a few large clones, younger patients have less curved LFIMs, as shown by the deviation from a straight line fit to the smallest 75% of clones (**Fig. 4**). Older individuals in contrast show a more distinct deviation from the line. This is consistent with simulations of the non-neutral competition (**Fig. 5**). At early timepoints, only the faster-growing non-neutral clones in the tail of the distribution are observed (**Fig. 5a**), leading to a straight line LFIM. A curve is observed at later timepoints once the slower-growing clones reach a size large enough to be detected (**Fig. 5b,c,d**). Future work with an increased number of patients is required to confirm this apparent trend.

**Figure 5.**
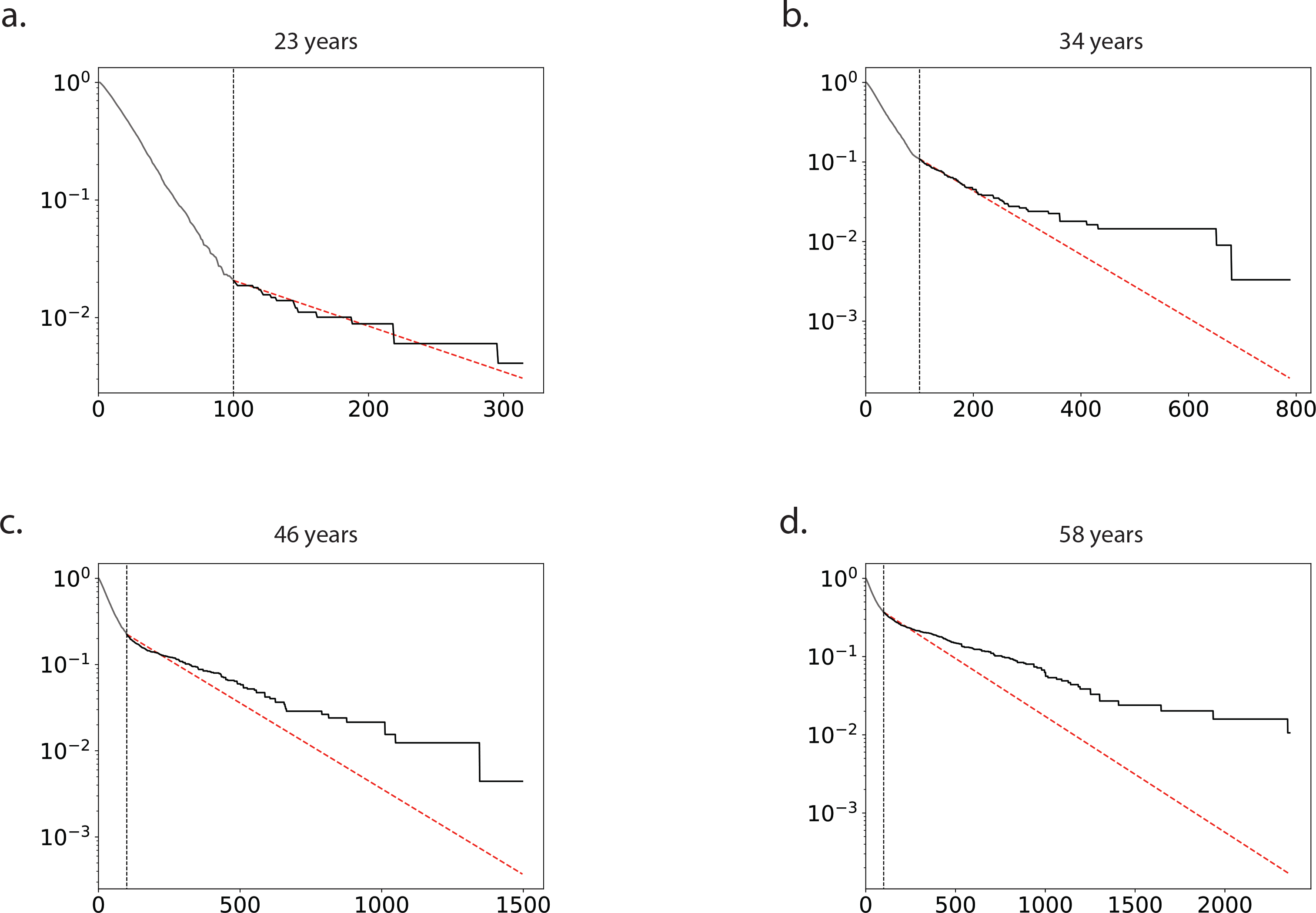
First incomplete moment over time. Curves in the LFIM may only be visible after sufficient time has passed, allowing both fast and slow growing clones to reach large enough sizes to be detectable through sequencing. Examples of the first incomplete moment for a simulation of non-neutral competition are shown for four timepoints. 1% of mutations are non-neutral with a fitness drawn from a normal distribution, N~N(mean = 0.1, std = 0.1). The vertical dashed line shows the detection limit, arbitrarily set at 100 cells. The section of the first incomplete moment that would not be visible due to the detection limit is shown in grey to the left of the line; the visible section is shown in black. The red dashed line is a straight line fit to the smallest 75% of *visible* mutant clones.

### Clone size as a marker of competitive selection of mutations

We have shown that clone size and LFIM alone cannot reliably classify clone sizes as neutral, due to a mixture of experimental limitations on the maximum and minimum sizes of clones and the fundamental effects of competition for space. In addition, where a curved LFIM is found, the position of the curve cannot simply discriminate the neutral and non-neutral mutant clones, although a trend of increasing proportions of non-neutral mutations at larger clone sizes is observed in both simulations (**Fig 6a,b**) and *in vivo* experiments (5). Neutral mutations hitchhiking on non-neutral clones may grow to large sizes, meaning that analysis restricted to synonymous mutations and mutations in non-expressed genes may not reflect purely neutral dynamics (**Fig 6c**). This raises a question of how to meaningfully interpret clone sizes observed in a tissue. This is an important question as there remain a small number of metrics for assessing the neutrality of mutations. While our results have demonstrated the risks of one specific interpretation of the LFIM, they also highlight the dangers of relying on a single measure of neutrality, especially if the underpinning mathematical assumptions are under-explored.

**Figure 6.**
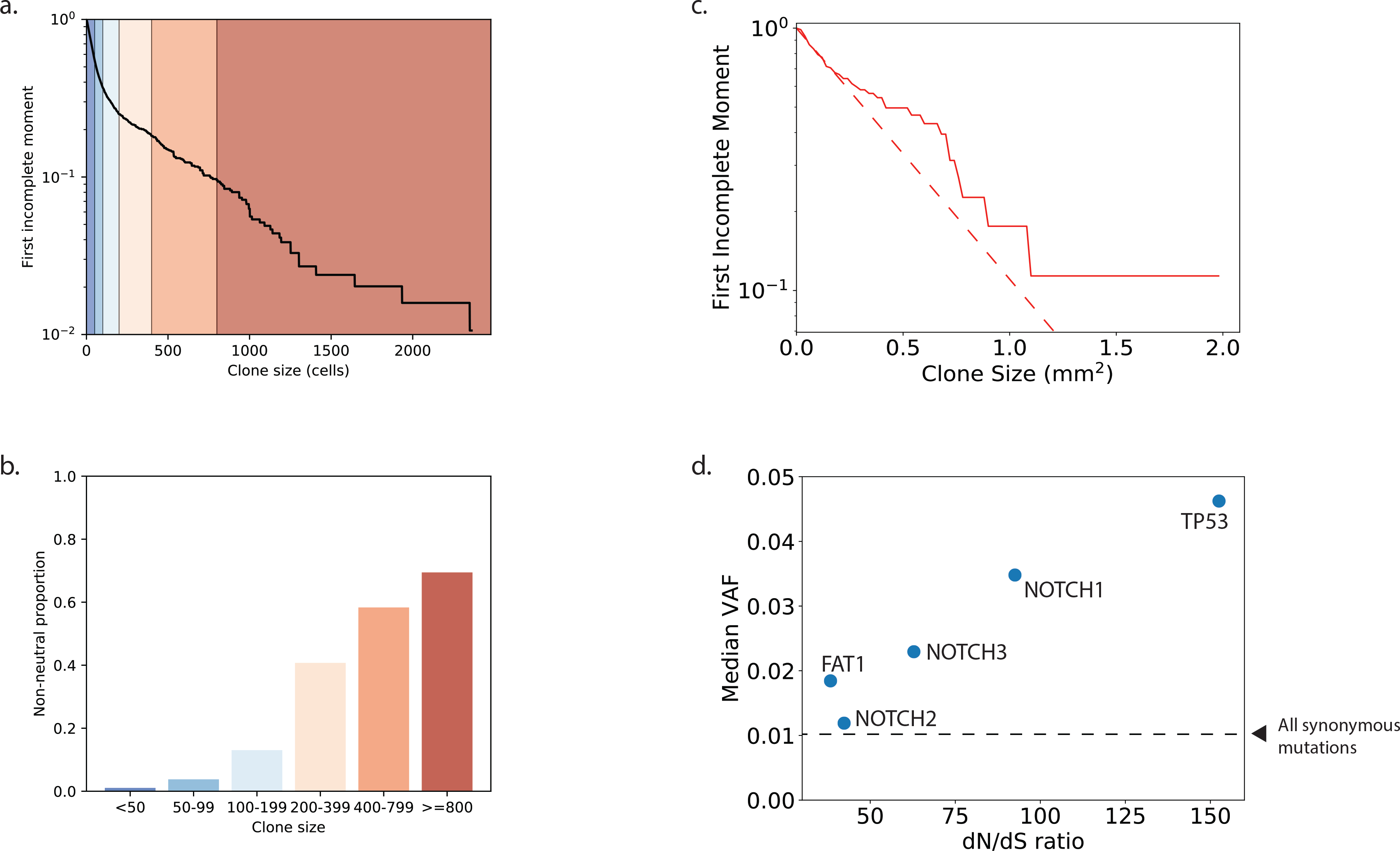
Clone size and selection. **a-b)** Proportion of non-neutral clones in different size ranges. **a)** The first incomplete moment of the clone size distribution from a simulation with 1% non-neutral mutations. Colored regions correspond to ranges of clone sizes described in **b**. **b)** Proportion of non-neutral clones in each clone size interval. Colors correspond to the regions shaded in **a**. **c)** First incomplete moments of the human esophagus mutation data for one individual, aged 72-75 (5), including only synonymous mutations and mutations in genes that are non-expressed. Non-expressed genes are selected using the Human Protein Atlas (35) (available from www.proteinatlas.org) using genes with a 0.0 TPM in the esophagus. The synonymous mutation T125T in *TP53* was excluded as it has been found to affect splicing (11, 26). Clone sizes which extend across multiple samples are merged using the methods of the original study (5). All mutations on chromosomes 1-22 were assumed to be heterozygous. The extent of deviation from the straight line can be seen by comparing the data (solid) to the dashed red line, which shows a straight-line fit to the smallest 75% of clones. **d)** Median variant allele fraction (VAF) for nonsense mutations in the five most significantly selected genes from the dN/dS analysis plotted against the dN/dS ratio for nonsense mutations. Combined results for all individuals in the study. The dashed line shows the median VAF of all synonymous mutations. Note that many of these synonymous mutations are likely to be passengers on non-neutral clonal expansions, and therefore the line does not represent the median VAF of mutations that have grown solely under neutral drift. One-sided Mann–Whitney tests show that, aside from NOTCH2 (p = 0.06), nonsense mutant clones in the genes shown are significantly larger than synonymous mutant clones (p-values < 0.0001).

In the specific case of the eyelid data, the original conclusion of non-neutral competition was supported by dN/dS analysis. While this is a widely used tool, this type of analysis is sensitive to the mutation model used for the neutral hypothesis (26), and detection of positive selection may be unreliable for some types of mutations in some genes. For example, almost all protein-truncating mutations inactivate the protein in which they occur. By contrast, missense mutations in some locations may reduce protein function, while in others they may generate a constitutively active mutant (30). In aggregate these effects may result in a dN/dS ratio close to 1.

Given the limitations of individual methods to assess neutrality, we speculated that combining discrete approaches may be more informative. To explore this we directly compared observed clone sizes and the associated dN/dS ratios of mutations in specific genes. We selected nonsense mutations from a panel of five mutated genes that were identified as under the strongest positive selection in normal esophagus and which have well-characterized roles in cancer (*TP53, NOTCH1, NOTCH2, NOTCH3*, and *FAT1*) (**Fig 6d**). For 4 genes there is the expected relationship between clone size and selection; that is, mutations in genes under greater selection pressure grow into larger clones. However, *NOTCH2* clones are under selection according to dN/dS criteria but have a similar size to synonymous clones.

There are multiple possible explanations for this unexpected result for *NOTCH2.* The dN/dS ratio indicates mutations that promote clonal expansion to a sufficient size to be detected. However, the impact of a mutation on clonal behavior may alter over time. This may occur if an initial expansion of a mutant clone increases the local cell density. If mutant cell proliferation is sensitive to this change of environment, the rate of clonal expansion may slow. Another potential mechanism is that the mutant clones grow initially due to an advantage over wild-type cells, but are later constrained by the growth of neighboring clones as the tissue is mutated over time. Both of these behaviors could lead to a high dN/dS ratio with only a modest increase in clone size, and are similar to observations of a *Trp53* missense mutation in mouse epidermis, where mutant clones have a strong competitive advantage over wild-type cells but their expansion is constrained (18).

The reverse observation, large clone sizes accompanied by only a modest dN/dS ratio, may indicate that mutations in a small region or hotspot in the gene can lead to extensive clonal expansion but mutations in the rest of the gene are under weaker selective pressure. An example from the esophagus data is *PIK3CA*, which has the largest median clone size of the 14 genes found to be under positive selection in human esophagus, largely due to the multiple large clones of the hotspot mutation H1047R. This highlights the importance of not just considering the gene in which a mutation occurs, but also the location of the mutation in the structure of the protein. Other factors such as epistatic interactions with other mutations (31) or age-related changes to the tissue microenvironment (32) could also lead to plastic and context-dependent mutant clone behavior and a complex relationship between dN/dS ratios and clone sizes.

## Discussion

We have presented two complementary explanations to resolve the apparent paradox regarding the dynamics of mutant clones in normal human eyelid skin. Both show how non-neutral competition can be consistent with a straight-line logarithm of the first incomplete moment of the inferred mutant clone size distribution – previously claimed to be an indication of neutral competition. Therefore, the mutant clone sizes observed in the normal human eyelid no longer appear to contradict the range of studies that suggest a number of mutations can drive non-neutral expansion of mutant clones in epithelia. We have also shown the benefits of using multiple orthogonal approaches to infer clone behavior. Finding a consensus can provide a high degree of confidence in the analytical conclusions, and inconsistencies may reveal an issue with one of the methods or help to identify interesting outliers in the data.

We have found that by considering spatial constraints of the tissue, non-neutral simulations can produce a straight-line LFIM, providing a counter example to the proposition that a straight-line LFIM implies neutral competition. We have also shown how the experimental method used to measure the sizes of mutant clones in the eyelid could have hidden signs of non-neutrality in the clone size distribution. Using isolated single samples which are too small in relation to the clone sizes will lead to underestimation of the size of a significant proportion of clones, as occurred in the eyelid experiment, and could lead to an apparently neutral clone size distribution. However, using over-large samples will reduce the ability to detect smaller clones, which can also lead to a straight-line LFIM because only the largest mutant clones are observed (23) (**Fig. 5a**). By using a grid of adjacent samples, the larger clones can be more accurately measured without compromising the detection of smaller clones, and can reveal the signs of non-neutrality that would otherwise have been hidden.

We have discussed the effects of sampling in detail. However, there are other potential confounding factors that could appear during DNA sequencing experiments. For example, comparing clone size distributions between genes may be hindered by variations in read coverage and the frequency of sequencing errors across genes, which could lead to different detection limits for small clones in different genes, and therefore different average clone sizes.

We conclude that mathematical and *in silico* models will be important tools for understanding clonal competition in pre-cancerous tissues. However, difficulties lie in the complex ways in which mutants grow, reacting to a changing local environment that may be altered by the mutant itself (18). Variation in mutation behavior may be seen both between and within genes, not to mention the complex ways in which mutants may interact with other mutants as neighbors or within the same clone. Exploring the consequences of adaptive mutant behavior, while still using models which are simple enough to fit to data and interpret, will be an ongoing challenge in the work ahead.

## Methods

### Simulations

The simulations were carried out on a 500 × 500 hexagonal lattice whose edges were wrapped to form a torus. Each cell was assigned a fitness value of 1 at the start of a simulation. Similar to a Moran process (27), one cell was randomly selected at each simulation step to differentiate (was removed from the simulation) and a neighboring cell was selected to divide to fill the space, with fitter cells having a higher chance of dividing (**Fig. 1f**). During each division, there was a chance that a mutation would occur in the new cell. If the cell did not mutate during division or if the mutation was neutral, the new cell produced would inherit the fitness of its parent cell. If the mutation was non-neutral, a random value drawn from a normal distribution, N~N(mean = 0.1, std = 0.1), was added to the fitness of a cell. We show in the **Supplementary Text** that our particular choice of distribution does not affect the conclusions of the analysis.

Estimates of cell cycle time in human tissues are hard to verify. However, we did not fit simulations to data, only demonstrated general properties of the models, hence the exact division rates used do not affect the conclusions of the analysis. For the neutral simulations, we used a division rate of 0.5 per week as estimated from the LFIM of clones in the human eyelid under the assumption of neutral competition (12). In the non-neutral simulations, the fittest clones can expand much faster than neutral, and therefore we reduced the division rate to 0.033 per week so that maximum mutant clone sizes were similar to the neutral simulations. The simulations ran for 3000 weeks (~58 years).

The somatic mutation rate for human tissue has been estimated at approximately 10^−9^ mutations per base pair per cell division (33) although we note that exposure to UV or mutagenic agents (such as stomach acid and alcohol) may substantially alter the mutation rate. With roughly 10^6^ base pairs included in the targeted sequencing experiment (4), this leads to a mutation rate of 10^−3^ mutations per cell division which we use for the neutral simulations. We use a higher mutation rate of 1.5 × 10^−2^ mutations per cell division for the non-neutral simulations so that the total clone numbers were similar in the neutral and non-neutral simulations.

Clone sizes were defined by the number of cells containing each mutation at the end of the simulation.

### Biopsy and sequencing simulations

Biopsies were simulated by taking 25 non-overlapping 70 × 70 cell squares from each grid. Assuming a density of basal cells of 10,000 per mm^2^ (34) and that half of basal cells are progenitors, this would make our biopsies approximately 1 mm^2^, similar in size to those used in the human eyelid (4).

Small clones may only appear in a very small proportion of sequenced DNA reads (if any) and are therefore hard to distinguish from sequencing errors (29), meaning they are not successfully detected as somatic mutations. To replicate this, we assumed a constant 1000x read depth and a requirement of 10 reads as a minimum to observe the mutant. For each mutant we had a true frequency *f*, the proportion of cells which contained the mutant. We assumed all mutations were heterozygous, so the true VAF was given by 0.5*f*. Each read then had a 0.5*f* chance of containing the mutation, so the total number of mutant reads observed, *reads*_*obs*_, was given by a draw from a binomial distribution with *n* = 1000, *p* = 0.5*f*. If *reads*_*obs*_ was greater than 10, we recorded the mutant as having a VAF of *reads*_*obs*_/*1000*, otherwise the mutant was unobserved and not included in the results.

### First incomplete moment test

We used the first incomplete moment as defined in (12).

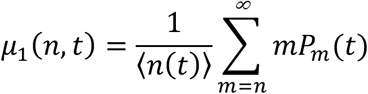

where *P*_*m*_(*t*) is the proportion of surviving clones that have *m* cells at time *t* and 〈*n*(*t*)〉 is the average mutant clone size at time *t*. As in previous studies (12, 23), we used R^2^, the coefficient of determination, to assess whether the log of the first incomplete moment was a straight line. The line fitting was constrained to pass through the point (*m*, 1), where *m* was the smallest observed clone size.

### dN/dS

Neutral and non-neutral mutations were introduced into the simulations with a known ratio, *a*. dN/dS was calculated as follows:

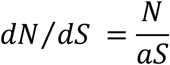

where *N* was the number of observed non-neutral clones and *S* was the number of observed neutral clones.

## Code Availability

Source code is accessible via Zenodo at DOI:10.5281/zenodo.1488100

## Acknowledgements

This work was supported by a Medical Research Council Grant-in-Aid to the MRC Cancer unit and core grants from the Wellcome Trust to the Wellcome Sanger Institute, 098051 and 206194. M.W.J.H. acknowledges support from the Harrison Watson Fund at Clare College. P.H.J. is supported by a Cancer Research UK Programme Grant (C609/A17257) and B.A.H. acknowledges support from the Royal Society (UF130039). We thank members of the Hall group at the MRC-Cancer Unit and the Jones and Martincorena groups at the Wellcome Sanger Institute for valuable discussions.

## Author Contributions

M.W.J.H, P.H.J. and B.A.H. wrote the paper. M.W.J.H wrote the code and ran the analyses. P.H.J. and B.A.H. supervised the study.

