## Supplementary figures and images for "Relating evolutionary selection and mutant clonal dynamics in normal epithelia"

### Supplementary Figure 1

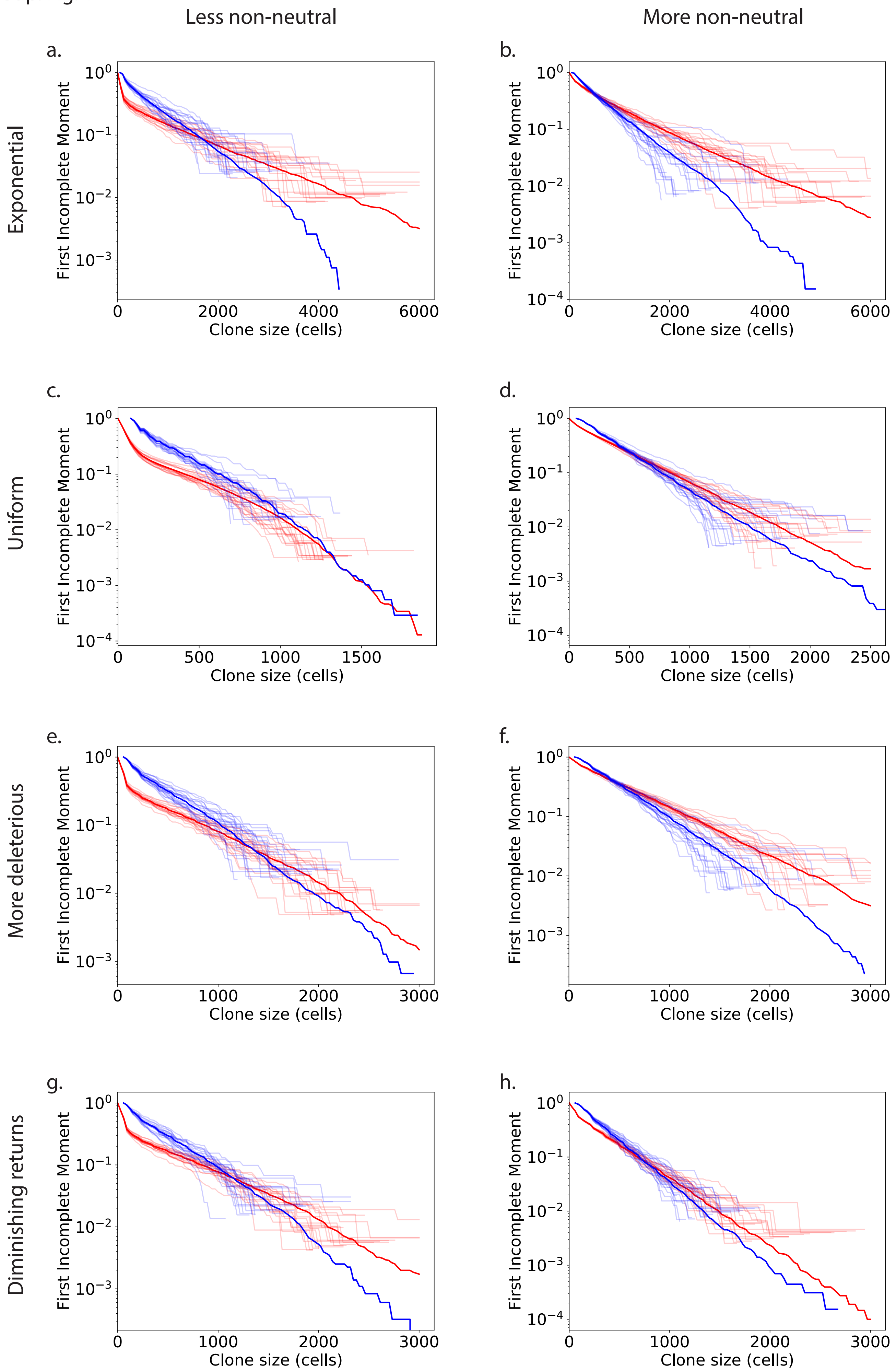
