## Supplementary Text for "Relating evolutionary selection and mutant clonal dynamics in normal epithelia"

### Distribution of fitness effects

The distribution of fitness effects (DFE) of the mutations can affect the shape of the LFIM of mutant clone sizes, and therefore the ability of the LFIM to detect non-neutral competition. This is shown by the difference between including 1% or 25% non-neutral mutations in simulations (**Fig. 2b,c and Fig. 3d**). Here we briefly explore how sensitive the shape of the LFIM is to the DFE of the non-neutral mutations, and therefore whether our conclusions were greatly influenced by the particular assumptions we have made. In particular we look at changing the shape of the non-neutral DFE, including additional deleterious mutations and changing the interaction of mutations (epistasis).

Several DFEs have been proposed based on theoretical predictions or experimental observations of mutations in evolving organisms (4). We have run simulations using three of the proposed distributions: normal (2, 3) (**Fig. 2b,c and Fig. 3b,c**), exponential (5) (**Sup. Fig. 1a,b**) and uniform (6) (**Sup. Fig. 1c,d**). In all of these cases the simulations produce similar shaped LFIMs.

In population genetics, it is generally assumed that a large majority of non-neutral mutations will be deleterious because an organism is already close to peak fitness (4). It is not clear to what extent this assumption applies for somatic mutations, since in healthy tissues individual cell fitness is secondary to the fitness of the organism as a whole (7). However, but it may still be the case that many non-synonymous mutations are deleterious to cell fitness. We have therefore run simulations in which two thirds of non-neutral mutations reduce the fitness of a cell, but this has little impact on the shape of the LFIM (**Sup. Fig. 1e,f**).

We have assumed in all cases so far that the effects of multiple mutations can be combined by simple addition of their fitness values. Here we look at diminishing returns (8) as an alternative form of epistasis. As a simple implementation of diminishing returns, we use the rule $new\ cell\ fitness = \mathrm{maximum}\ (old\ cell\ fitness,\ new\ mutation\ fitness)$, i.e. the fitness of a mutation will replace (rather than add to) the cell fitness if it is higher than the cell fitness, otherwise it has no effect. This rule means that a very fit clone will only rarely increase in fitness through a new mutation and the change is likely to be small. The results of these simulations are shown in **Supplementary Figure 1g, h** and are similar to those using simple addition of mutation fitness.

The true effects of somatic mutations are likely to be far more complex than the examples we have explored here (1). However, in all cases we have simulated we see that the two key observations are not altered: Firstly, the use of biopsies reduces the curve in the LFIM and therefore reduces the ability of the LFIM to detect non-neutral competition. And secondly, a straight line LFIM does not necessarily imply non-neutral competition, as shown in the highly non-neutral simulations.

**Figure Legend**

**Supplementary Figure 1 -** ***LFIM for alternate distributions of fitness effects***

For all cases, the LFIMs using full clone sizes is shown in red and the LFIMs using clone sizes observed through simulated biopsies and sequencing is shown in blue. 20 individual simulations are shown for each case and the mean of 100 simulations is shown in bold. Except where stated otherwise, the simulations in the left column are 1% non-neutral, and the simulations in the right column are 25% non-neutral. All simulations are run on 500 x 500 grids, have a division rate of 0.033 per week, a mutation rate of 0.015 per cell division and are run for 3000 weeks (~58 years). Unless stated otherwise, the cell fitness is the sum of the fitness of all mutations in the cell.

**a,b)** The fitness of non-neutral mutations is drawn from an exponential distribution with parameter 0.1. **a)** 1% of mutations are non-neutral. **b)** 25% of mutations are non-neutral.

**c,d)** The fitness of new mutations is drawn from a uniform distribution, U~(1, 1.2). **c)** 1% of mutations are non-neutral. **d)** 25% of mutations are non-neutral.

**e,f)** Addition deleterious mutations. **e)** The fitness of 1% of new mutations is drawn from N~(mean = 0.1, std = 0.1) (mostly beneficial mutations) and the fitness of another 2% of new mutations is drawn from N~(mean = -0.3, std = 0.1) (mostly deleterious mutations). The remaining 97% of mutations have no effect on cell fitness. **f)** The fitness of 25% of new mutations is drawn from N~(mean = 0.1, std = 0.1) (mostly beneficial mutations) and the fitness of another 50% of new mutations is drawn from N~(mean = -0.3, std = 0.1) (mostly deleterious mutations). The remaining 25% of mutations have no effect on cell fitness.

**g,h)** Diminishing returns. The fitness of new mutations is drawn from N~(mean=1.1, std=0.1). $New\ cell\ fitness = \max\,(old\ cell\ fitness,\ \ new\ mutation\ fitness)$, i.e. the fitness of a mutation will replace (rather than add to) the cell fitness if it is higher than the cell fitness, otherwise it has no effect. **g)** 1% of mutations are non-neutral. **h)** 25% of mutations are non-neutral.

Sup. Fig. 1

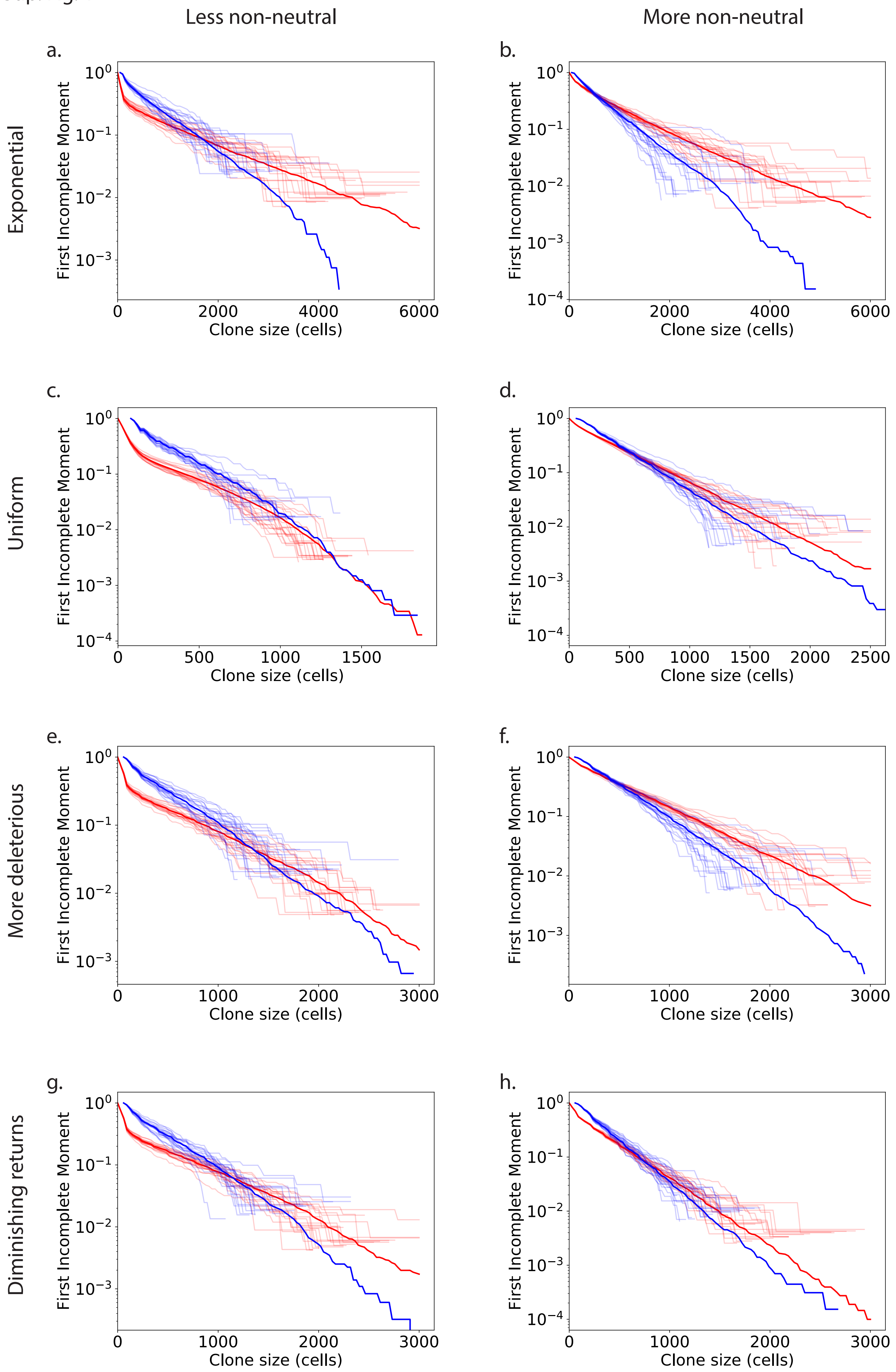
Less non-neutral
More non-neutral
Exponential
Uniform
More deleterious
Diminishing returns
a.
b.
c.
d.
e.
f.
g.
h.
First Incomplete Moment
Clone size (cells)
$10^0$
$10^{-1}$
$10^{-2}$
$10^{-3}$
$10^{-4}$
0
500
1000
1500
2000
2500
3000
4000
6000
